# DigiFish: A flexible open-source tool for animating realistic virtual fish and reconstructing visual fields from tracking data

**DOI:** 10.64898/2026.10.02.755892

**Authors:** Katie Dunkley, Jolyon Troscianko, Anna Ahlberg, Lauren Neville, Christos C. Ioannou, Robert J.P. Heathcote

## Abstract

1. For many animals, vision represents the primary sensory modality governing a vast diversity of behaviours. Creating ‘virtual twins’, or visually realistic digital reconstructions of real individuals interacting within their environments, offers a promising approach for understanding how visual information guides an animal’s decisions, behaviours, and interactions. Such systems enable the creation of video scenes from any view, including the viewpoint of the animals themselves. This allows receiver information to then be analysed from the perspective of species-specific visual systems, or for the accurate reconstruction of a real animal’s movement to provide social stimuli for behavioural experiments.
2. Despite advances in 3D modelling, photorealistic scene rendering, and animal tracking, efficiently creating realistic digital reconstructions of movement remains challenging for many species. In fishes, a continuously-deforming body due to the absence of discrete joints make it particularly difficult to map tracked body keypoints onto articulated digital models, often resulting in unnatural or erratic animations.
3. Here, we introduce DigiFish, an add-on for the open-source animating software Blender, that generates realistic fish animations from tracking data. By integrating anatomical keypoint tracking with spline-based body curvatures, DigiFish reconstructs biologically-realistic swimming postures and movements of fishes within customisable virtual aquatic environments. The system also supports virtual ‘point-of-view’ cameras that can be placed arbitrarily within a scene, or dynamically driven by eye-tracking data, enabling unlimited perspectives on how individuals, conspecifics, or landmarks appear against complex backgrounds and how these views change with realistic swimming and eye movements.
4. We provide the Blender add-on and user guide and use DigiFish to reconstruct two different contexts: staged predator-prey interactions in the laboratory and natural schooling of wild fish in the field. Using a further reconstruction of fish swimming in an arena, we also use an Elementary Motion Detector model to show that our virtual reconstructions successfully capture perceptually-realistic movements and postures of individual fish within a digitally reconstructed scene.
5. By integrating photo-realistic scenes with biologically-realistic, simulated motion, DigiFish offers a flexible and accessible platform for investigating visually guided behaviour, perception, and cognition in complex and dynamic environments.

## 1 Introduction

Each animal experiences the world through its own *Umwelt*, a sensory reality determined by the information it extracts and processes from its environment (Von Uexküll, 1957). Among the sensory modalities contributing to an animal’s *Umwelt*, vision is often the dominant source of information, guiding an animal’s decisions, behaviours, and interactions. However, a central challenge in behavioural ecology has been accurately modelling how this visual information is perceived given varying constraints across animal sensory processing systems (Cronin et al., 2014; Endler, 1992; Stevens, 2013). While advances in comparative perceptual psychology now allow us to model how visual information may be received and processed by different species, such as via models of visual acuity, motion detection, and spectral and contrast sensitivity (e.g. Borst & Egelhaaf, 1989; Caves et al., 2025; Caves & Johnsen, 2018; Endler & Mielke, 2005; Kelley & Endler, 2017; Santon et al., 2025; Troscianko & Stevens, 2015), these approaches still require realistic representations of the visual information an animal actually obtains as it moves through its environment. Obtaining such observer-specific, point-of-view footage remains an important challenge, particularly given how even small deviations in viewing position can result in fundamentally different information obtained by an observer (Hogan & Stoddard, 2018; Kelley & Endler, 2017; Li et al., 2026). Although animal-borne cameras can help reconstruct an individual’s viewpoint (Moll et al., 2007), their utility is mostly confined to relatively large species. One alternative approach is to create animated ‘virtual twin’ versions of real animals as they interact within their environment (Bian et al., 2021; Iwata et al., 2026; Miñano et al., 2023; Wallace et al., 2025; Wu et al., 2022, 2024). This approach allows realistic point-of-view videos to be rendered from anywhere within the scene, including that of the animal’s own hypothetical visual field, offering enormous flexibility to address how visual information can affect behaviour. Such digital scenes go beyond traditional animated stimuli (e.g. Chouinard-Thuly et al., 2017), and, following conversion and calibration of videos via suitable visual modelling approaches, provide us a step closer to seeing as animals see.

Following advances in computational power and machine learning techniques, the generation of visually-realistic 3D models of animals (Irschick et al., 2022; Miller et al., 2022) and their natural environments (Yang et al., 2025; Zhang & Johnson-Roberson, 2023, 2023) are making the construction of such virtual twins increasingly accessible. Coupled with automated marker-less video tracking of animal body parts (for review see Mathis et al., 2020), naturalistic postures and movement trajectories can be mapped onto these digital models with high spatiotemporal fidelity (Lobato-Rios et al., 2022; Luo et al., 2022; Plum et al., 2023; Vaxenburg et al., 2025; Wu et al., 2022, 2024). Together, these developments mean that dynamic reconstructions of real animal interactions can increasingly be generated without prohibitively specialised resources (e.g. Wallace et al., 2025).

Despite these advances, applications of these methods have understandably focused on animals with relatively rigid bodies and identifiable and easily-tracked inter-joint angles and fixed bone lengths (e.g. insects: Plum et al., 2023; Vaxenburg et al., 2025). However, many animals have bodies that bend and deform continuously. Fishes, for example, which account for around half of all extant jawed vertebrate species (Gumbs et al., 2024), lack distinct joint locations along their body, and their body shape can be difficult to accurately reconstruct onto an articulated 3D model from tracked keypoints. As such, posture and movement reconstructions are prone to erratic, distorted outputs that fail to capture the smooth, undulating nature of these animals’ movements. Recent work has demonstrated that realistic fish motion can be digitally recreated using machine learning approaches such as image segmentation (Wu et al., 2022), showing this problem is technically tractable. However, these approaches can be difficult to apply in practice due to their substantial computational requirements and need for specialist expertise.

Here we present DigiFish, a free, open source, user-friendly, Blender add-on for reconstructing visually-realistic digital fish and their behaviour from real-animal tracking data. DigiFish can be used to construct and model visual scenes from any receiver’s perspective, or to generate naturalistic experimental stimuli based on realistic movement of real animals, that can subsequently be readily manipulated and presented to live animals using digital displays (e.g. Ingley et al., 2015). Blender is a powerful open-source 3D modelling and animation platform that supports the entire animation workflow, from physical modelling and rigging, to animation and rendering (Community, 2018; Flavell, 2011). It is already widely used in biological and ecological research to produce anatomically accurate models (Hancock et al., 2026; Khan et al., 2019) and explore visually realistic environmental scenes (e.g. Hillen-Schiller et al., 2025). DigiFish integrates with Blender’s standard animation workflows and gives users full control over body postures and swimming movements of single or multiple fish(es) within fully customisable scenes (e.g. background, lighting, water clarity). To capture the natural undulating and oscillating motions of swimming fish, DigiFish employs Bezier curve–based rigging, avoiding the constraints of fixed keypoint distances or rigid skeletons and allowing smooth, lifelike movement across the whole body of the fish. Critically, virtual ‘point-of-view’ cameras can be fixed or dynamically positioned (using eye-tracking data from the real fish) on digital fish, allowing us to render videos that can approximate the dynamic visual field of a swimming fish. By combining realistic appearance with flexible, natural movement, DigiFish opens new opportunities for investigating visual perception, social behaviour, and predator-prey interactions. Here, we provide a walkthrough of the DigiFish add-on and demonstrate its use in both lab and field environments.

## 2 DigiFish Blender Add-on Overview

DigiFish’s simple graphical user interface has been designed to guide users through the full steps of the animation and rendering process for single or multiple fish(es) (Figure 1). In addition to the add-on, we provide detailed step-by-step instructions on the project’s GitHub (https://github.com/katie-dunkley/DigiFish). DigiFish was developed with accessibility at the forefront so requires no coding experience and is applicable across a variety of biological fields (see *Applications*). Below we briefly outline the main steps of this pipeline.

**Figure 1:**
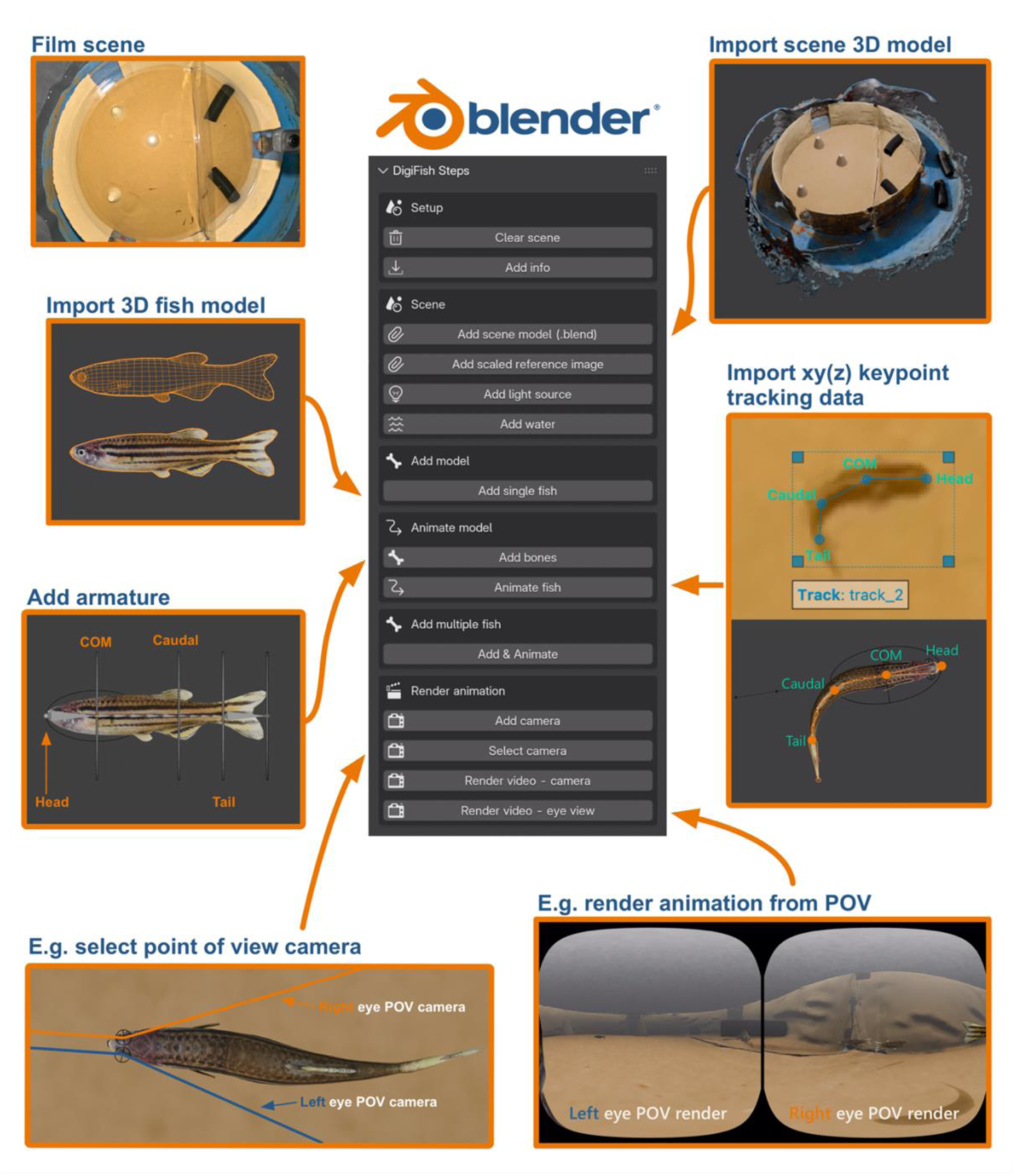
The steps to add and animate digital fish with DigiFish. Examples shown are from the lab reconstruction (see *Table 1* for information and model credits, POV: point-of-view, COM: centre of mass).

**Table 1:** Outline of setups used for testing DigiFish in both a laboratory and field environment.

| DigiFish Steps | Lab | Field |
| --- | --- | --- |
| Species | 4 × zebrafish (*Danio rerio*) and 2 × tigerfish predator (*Datnioides microlepis*) | 3 × Eurasian minnow (*Phoxinus phoxinus*) |
| Location | John Krebs Field Station, University of Oxford, Wytham, UK | Letcombe Brook, East Hanney, Oxfordshire, UK |
| Environment | 80 cm diameter arena with sandy bottom and acetate partition separating predator and prey. | Shallow chalk stream with rocky/stone substrate. Turbid water. |
| 3D model generation | Zebrafish: sculpted in Blender with UV mapping by Andreas Øium.<br>Tigerfish predator: purchased from TurboSquid | Minnow: sculpted by K.D. in Blender with UV mapping |
| 3D scene generation | Reconstructed using Luma AI (https://lumalabs.ai/) and exported as .obj file (filmed with GoPro Hero 13, 2K resolution). | Reconstructed using photogrammetry and Metashape Pro (filmed with Insta360 x4, 5.7K resolution). |
| Camera setup | Overhead view: GoPro Hero 13, 240 fps, 2.7K resolution, linear field-of-view.<br>Lateral view: GoPro Hero 13, 240 fps, 2.7K resolution, linear field-of-view<br>Camera for predator eye tracking (second dorsal view): Insta360 x4, 30 fps, 8K resolution (tracked in 4K) | Dorsal view: GoPro Hero 13, 240 fps, 2.7K resolution, linear field-of-view<br>Lateral view: GoPro Hero 13, 240 fps, 2.7K resolution, linear field-of-view |
| Lighting | NEEWER MS150B 130W Bi-Color LED Video Light setup directly overhead | Natural overhead sunlight |
| Tracking software | SLEAP.ai (Pereira et al., 2022) used on overhead view footage |  |
| Keypoints | Zebrafish: 4 x (shown in Figure 1)<br>Predator: 6 x (+ 4 x points for eyes, shown in Figure 2) | Minnow: 4 x |

### 2.1 Animal three-dimensional model

DigiFish prompts users to import a blend file containing a 3D model of the fish species they wish to animate. These digital models can be created using a variety of methods, including photogrammetry (e.g. Álvaro-Tordesillas et al., 2023; Bot et al., 2019; Miller et al., 2022), AI-generated meshes (e.g. https://aidemos.meta.com/segment-anything, https://www.meshy.ai), or Blender’s own sculpting tools (e.g. Flavell, 2011). Users can also import pre-made models from online marketplaces (e.g. https://www.turbosquid.com/). DigiFish appends the blend model file directly to the existing project to preserve the original materials, textures, and shading information of the model (irrespective of the method used for model generation), helping to simplify the animation pipeline for DigiFish users.

Imported models should include any relevant skin colouration, created either by mapping spectrally calibrated or reference photographs onto the model surface using Blender’s UV mapping tools, or by applying colour directly through Blender Dynamic Surface Painting (BDSP), which enables surface-level painting and editing (Flavell, 2011). Users can also simulate fine-scale fish texture features like fish scales or silvery, glossy colouration via BDSP, although such detail may be unnecessary for most applications given the markedly increased rendering time.

If needed, species-specific visual modelling can be incorporated at different stages of the DigiFish workflow depending on the intended application such as visual analysis, or presentation of videos to real animals (the latter may also require appropriate calibration of the display device). For instance, modelling the perceived spatial resolution of the scene must be performed after video rendering, whereas chromatic modelling based on animal cone catches can be incorporated either before or after (e.g. with the MICA Toolbox; Troscianko & Stevens, 2015 or AcuityView; Caves & Johnsen, 2018). Cone-catch-calibrated images, for example, can be directly used as input images to generate the textures for 3D fish models, or the visual scene can be modelled post-rendering when appropriate colour standards are incorporated into the digital environment.

### 2.2 Animating with DigiFish

Through DigiFish’s user interface and keypoint tracking data (Figure 1), users can animate their model fish. DigiFish uses keypoint tracking data, provided in either two (x-y) or three (x-y-z) dimensions, along with a pixel-to-centimetre conversion value, to animate changes in fish body posture and position through time and space.

#### Keypoints and tracking data

DigiFish requires two user inputs: (1) metadata specifying the names and proportional locations of each anatomical keypoint along the fish’s body (as used in the tracking), and (2) a CSV files containing frame-by-frame tracking data of the keypoints. Tracking data can be generated from collected videos using open-source tools such as DeepLabCut or SLEAP.ai (Mathis et al., 2018; Pereira et al., 2022), and 3D tracking is possible using stereo setups (e.g. Dunkley et al., 2023). For accurate reconstructions of postures during swimming, we recommend overhead cameras with sufficiently high frame rates to generate keypoints at a frequency of at least twice the tail-beat frequency. DigiFish will then interpolate missing x-y(-z) coordinates across untracked frames via smoothing (see below). Users can also upload simulated tracking data (i.e. if using DigiFish for stimulus presentations): fish posture data for example can be generated by cycling through angular changes between consecutive keypoints along the body (e.g. see Faria et al., 2019 for detail).

We recommend users define at least four keypoints along the fish’s dorsal body axis (e.g. Figure 1, Table 1). The two most anterior points, usually head and centre of mass (species dependant), maintain a straight line regardless of body curvature (as also in Qian & Chen, 2017) and are used to determine fish yaw, pitch, and spatial position (see “Orientation” and “Location” below). These two points alone can animate rigid models, but accurate body curvature, swimming motion, and dynamic posturing require at least four keypoints in total. Users should ensure the number and placement of keypoints capture sufficient spatial resolution for accurate curvature estimation depending on the morphometrics of their study species (see “Posture”). Because transparent tails are difficult to track and may provide little visual information, DigiFish can estimate caudal fin position based on body curvature if this keypoint is omitted.

#### Posture

DigiFish reconstructs posture using keypoint coordinates, relative angles (as in Faria et al., 2019), and a natural cubic spline interpolation passing through all keypoints (highlighted in Martineau & Mourrain, 2013). It maps user-supplied keypoints (x-y from overhead view) to Inverse Kinematics (‘IK’) controllers along the model fish’s armature (digital proxies for bones) in Blender, which subsequently controls a Bézier curve that deforms/bends the fish mesh (Figure 1), constraining body bending to the most anatomically plausible patterns. Users can adjust body deformation smoothing by varying the number of armature segments between keypoints which simulate individual vertebrae. To correct for tracking noise caused by fishes’ lack of discrete landmarks from overhead footage (Zheng et al., 2024), DigiFish applies LOESS smoothing independently to each keypoint’s x and y coordinates across frames (users can adjust smoothing through the GUI). DigiFish then fits a smoothed curve through the keypoints, rescaling their positions to ensure accurate inter-keypoint distances relative to the straight reference model. Missing terminal keypoints (e.g. end of tail) are estimated from the curve shape. Smoothing and mapping are performed automatically.

#### Orientation

The yaw (horizontal orientation) of the model fish is determined by the bearing of a straight line connecting the first two keypoints (e.g. the head/nose and centre of mass, Figure 1). After applying LOESS smoothing, DigiFish calculates how much an undulating fish changes its overall bearing across frames (the ‘winding angle’, Siciliano & Khatib, 2016). This angle controls rotation around the second most anterior keypoint, enabling smooth and realistic yaw transitions: turning right increases the cumulative angle, turning left decreases it. DigiFish will also animate body pitch (head-up/down posture, see GitHub for instructions) if the data are supplied by the user (pitch angle, with negative values indicating head-down posture). For the many applications where 2D tracking is sufficient (Miller & Gerlai, 2012; Romenskyy et al., 2020) only yaw is animated, with pitch fixed at 0°.

#### Location

DigiFish reconstructs movement based on the x–y (and optionally z) coordinates of the second user-defined keypoint, which are converted to real-world units once users supply the required pixel-conversion scaling factor. DigiFish applies LOESS smoothing to the coordinate values of this second keypoint across frames to produce smooth, continuous motion (users can adjust the degree of this smoothing through the GUI). Users should ensure their imported tracking data uses the same bottom-left origin coordinate system (0,0) as Blender; many tracking tools default to top-left, which will cause misaligned animation within the scene.

### 2.3 Scene reconstruction

As part of the DigiFish workflow, users can import and embed their animated fish within realistic 3D environmental scenes, allowing researchers to use rendered videos as biomimetic stimuli and/or test how fish perceive and interact with their environment.

#### Scene

Recent advances in image-based 3D reconstruction now allow users to create high-quality digital environments with relative ease. DigiFish supports scenes generated using multiple methods, such as structure-from-motion photogrammetry (e.g. Bayley & Mogg, 2020), Neural Radiance Fields (‘NeRFs’; Zhang & Johnson-Roberson, 2023), and Gaussian Splatting (Yang et al., 2025, imported using a further add-on e.g. <u>Kiri Innovation add-on</u>, <u>ReshotAI add-on</u> see <u>ReshotAI guide</u>). NeRFs use neural networks to reconstruct photorealistic 3D scenes from 2D images, while Gaussian Splatting offers a faster, less computationally intensive alternative using 3D Gaussian functions for real-time rendering and interactive scene manipulation (Chen & Wang, 2024). These reconstruction techniques require no specialist equipment (e.g. Cerbone et al., 2025) and can be accessed via free platforms like Luma AI (https://lumalabs.ai/) or KIRI Engine (https://www.kiriengine.app/). DigiFish works by appending the user’s .blend scene file directly into the animation project, preserving materials and properties for use with the animated fish(es). To assist with scaling and alignment, users are encouraged to import a reference image (e.g. a video screengrab) to use as a guide and use a scale bar within the scene.

#### Water

Water, and its dissolved and suspended contents, can alter the bending, scattering and absorption of light (Kitchener et al., 2017; Utne-Palm, 2002). As such users may want to incorporate these visual effects into their scene, and DigiFish begins to help users to do this by adding an optional, computationally-efficient fog-like volumetric material that still facilitates fast rendering. Users can modify the Principled Volume settings (e.g. colour, density, directional light scattering (‘anisotropy’)) of the water material via a simple user interface that appears when the water object is selected. Through DigiFish, users can also animate water movement within the scene to simulate how water current affects turbidity movement. DigiFish was developed primarily to replicate fish movement and body position from tracking data and uses a default computationally-efficient rendering method.

Therefore, it does not explicitly attempt to accurately model the optical effects of water under more complex conditions, but there are many additional realistic ways to model water in Blender (e.g. with volumetric raytracing, see Nakath et al., 2022, or with Blender’s Fluid Simulator, Mullen, 2008) and these can be incorporated alongside DigiFish if needed by the user.

#### Lighting

DigiFish adds a configurable digital light source (‘area light’) to a project and provides a user interface to adjust the light’s (1) colour, (2) location, (3) intensity (power in watts), and (4) whether it casts shadows. Users should fine-tune these lighting settings in combination with water properties (see above) to visually match the real environment. For sensory ecology projects requiring more accurate or spectrally-calibrated reconstructions not included in DigiFish, such as those involving visual modelling, users can incorporate Blender’s advanced lighting techniques (e.g. ray tracing), which accurately simulates how light interacts with objects (e.g. refraction/absorption), or caustic lighting effects (e.g. Iraci, 2013). However, these techniques demand significantly more computational power for rendering animations and may be unnecessary for most DigiFish applications.

### 2.4 Rendering

DigiFish supports rendering from two viewpoint types: a simulated fish’s perspective, or user-defined perspectives positioned anywhere within the scene. First, for fish-eye views, virtual cameras are fixed at the fish’s left and/or right eye (Figure 1). To mimic the wide visual field of many fish (rather than the standard narrow-angle cameras in Blender), DigiFish uses the eeVR add-on (https://github.com/EternalTrail/eeVR) for panoramic rendering. Users can set the total field of view (e.g. up to 180°) based on species-specific visual information. Importantly, eye cameras can also move with supplied eye-gaze tracking data, which captures the eye vector’s posterior-to-anterior orientation relative to the first two body keypoints: negative angles indicate that the eye faces inwards towards the snout, and positive angles outward towards the tail (Figure 2, see code on project GitHub). DigiFish assigns rotations to the eye cameras, allowing researchers to explore how a fish’s visual field shifts with body and eye movements and to render the scene from the fish’s own perspective. This creates new opportunities to investigate how fish perceive conspecifics, predators, and other features of their environment while moving.

**Figure 2:**
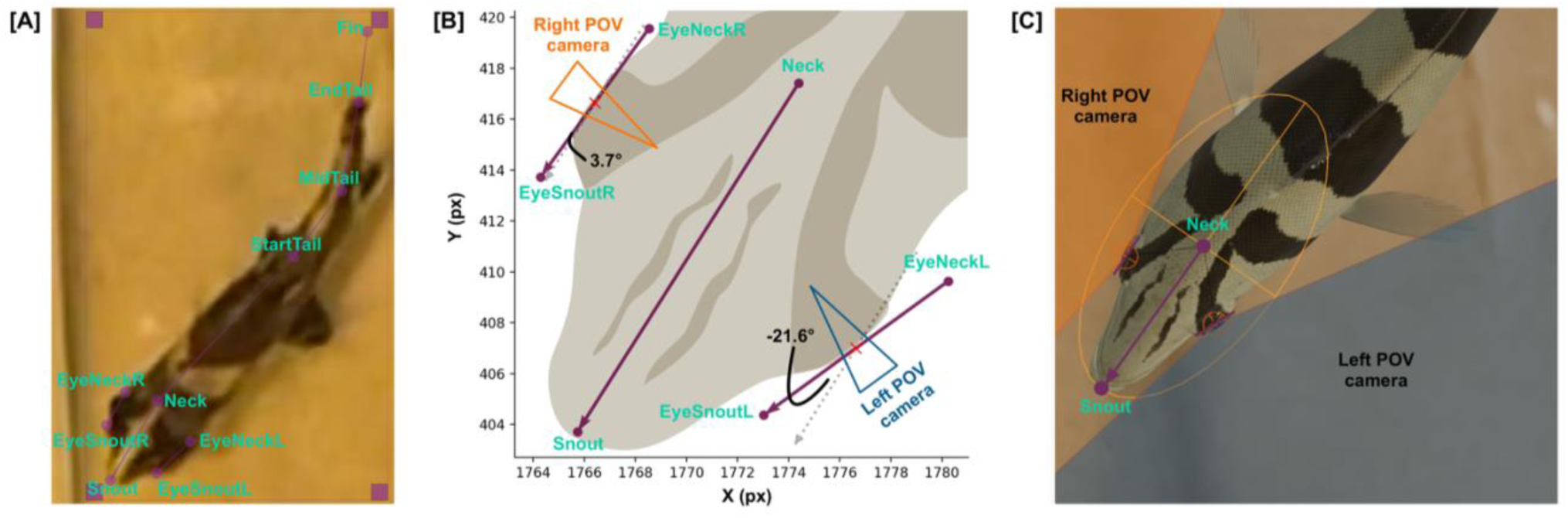
Method for calculating eye orientation angles used to dynamically orient eye point-of-view cameras in DigiFish. (A) SLEAP.ai tracking keypoints used to define the body and eye vectors for Datnioides microlepis. Eye orientation is represented by the vector from ‘EyeNeck’ to ‘EyeSnout’ for each eye. (B) Eye orientation is defined as the angle between each eye vector (purple line) and the vector perpendicular to the body midline (defined by the Neck–Snout vector, parallel vectors for each eye shown by grey dashes). This angle is then used to rotate the corresponding eye direction control wireframe in Blender. Negative angles indicate that the eye is rotated inward toward the snout, whereas positive angles indicate that the eye is rotated outward toward the tail. (C) The rotated control wireframes determine the orientation of the dynamically moving point-of-view cameras in DigiFish. For fishes where these data are not available, point-of-view cameras remain fixed pointing outwards (e.g. see zebrafish in Figure 1).

Second, for user-defined visual perspectives, DigiFish allows virtual camera placement anywhere in a scene. These views can be used to assess animation realism (e.g. through comparison with real video footage; see DigiFish Testing), visual modelling of hypothetical observers (e.g. a predator), or to generate visually realistic and flexible social/behavioural stimuli of virtual fish (with or without backgrounds) for behavioural experiments. Users can also modify these camera views to simulate flat or dome underwater camera ports (e.g. Menna et al., 2024; Nakath et al., 2022) but this functionality is not included in DigiFish itself.

## 3 DigiFish Testing

We trialled DigiFish under two different contexts (see Table 1 for detailed steps): staged predator-prey interactions between Indonesian tigerfish (*Datnioides microlepis*) and zebrafish (*Danio rerio*) in the lab (Figure 1), and natural schooling of wild European minnows (*Phoxinus phoxinus*) in a British chalk stream (lab: https://tinyurl.com/DigiFishLab, field: https://tinyurl.com/DigiFish).

To assess the realism of our DigiFish simulations, we used an Elementary Motion Detector model (Hancock et al., 2026; Santon et al., 2025) (n = 8) parameterised to fish spatial and temporal visual frequencies to compare luminance-based visually relevant motion signals from our simulations against those from real fish video footage (see Supplementary Materials for method). As illustrated in Figure 3, our virtual reconstructions successfully capture realistic movements and postures of individual fish, based on a biologically-informed model of motion detection, within a digitally reconstructed scene.

**Figure 3:**
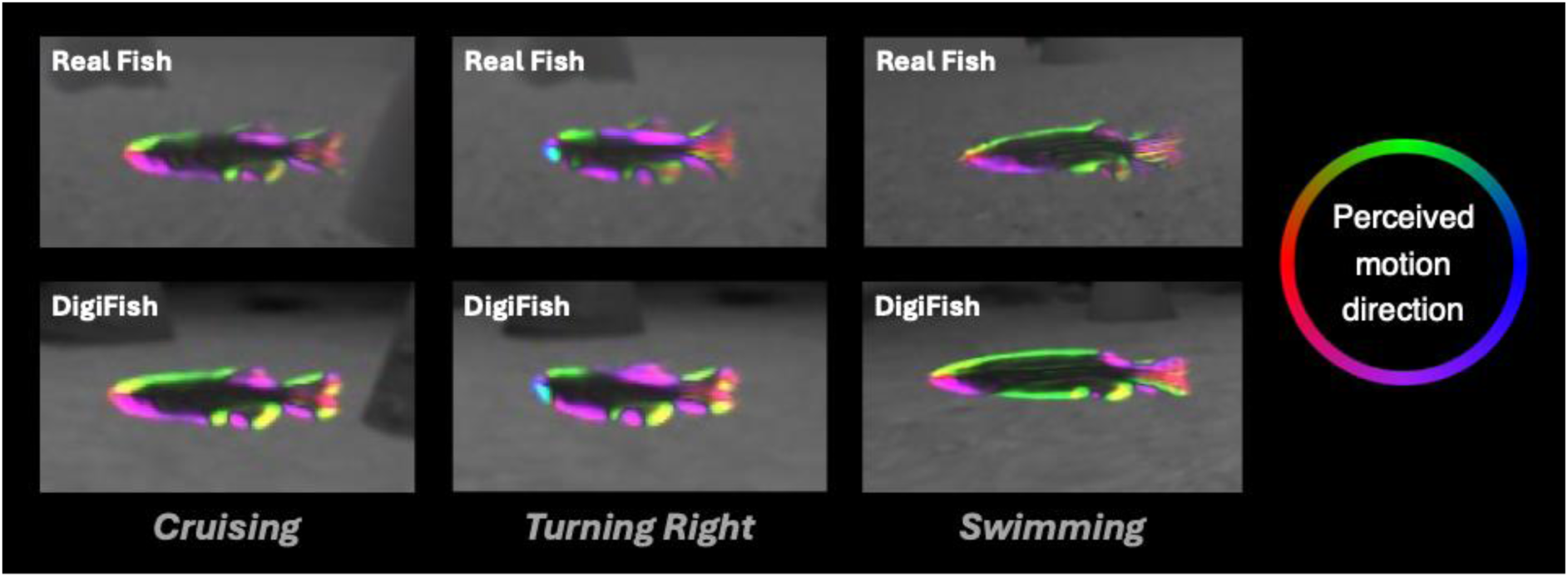
Colour output from an elementary motion detector applied to real versus rendered footage showing how movement is perceived in the virtual reconstruction compared to the real scene (for methods see Supplementary Materials).

## 4 DigiFish Applications & Conclusion

By recreating realistic 3D postures and movements of fish(es) within controlled visual scenes, DigiFish provides a powerful, tractable tool for understanding how fish move, perceive, communicate, and interact within their visual environments. By placing virtual cameras that move with virtual fish or sit anywhere within a scene, DigiFish offers near-infinite perspectives on how individuals, conspecifics, or landmarks appear against complex backgrounds, and how these views change dynamically with realistic swimming and eye movements. Beyond this observational framework, Blender and DigiFish’s open-source flexibility lets users manipulate the appearance and morphology of virtual fish (e.g. colour patterning, body shape), as well as the structure, visibility, and composition of their habitats. Such control enables users to generate lifelike visual stimuli, alter or isolate specific cues, and create reproducible variants of the same interaction, forming a powerful experimental framework for fish behavioural studies. Renders from DigiFish can thus be used to address a broad range of questions centred on how visual information shapes behaviour, including within- and between-species interactions, communication, navigation, foraging, and decision-making. These rendered scenes can also be presented to live individuals as stimuli through physical displays (e.g. Chouinard-Thuly et al., 2017) or further developed as virtual reality-style digital twins (e.g. Li et al., 2025; Sanchez et al., 2025; Stowers et al., 2017), enabling tightly controlled, repeatable behavioural assays within fully manipulable visual environments. For example, visual scenes can be rendered at the high frame-rates that are critical for accurately modelling visual motion that is free from perceptual artefacts (e.g. Hancock et al., 2026).

As automated tracking and 3D scene-generation methods continue to improve, the strength and realism of DigiFish will only grow. Its flexible, easy-to-use, and open-source design, combined with reconstructions grounded in true biological movement, positions DigiFish as a practical platform for next-generation research on visually guided behaviour, fish perception, and cognition in complex and dynamic environments. Together, these features establish DigiFish as a unique and timely resource for the study of fish in their visual worlds.

## Author contributions

**K.D.:** Conceptualization, Data curation, Formal analysis, Investigation, Methodology, Project administration, Software, Supervision, Validation, Visualization, Writing – original draft**. J.T.:** Conceptualization, Funding acquisition, Methodology, Resources, Writing – review & editing**. A.A.:** Formal analysis, Investigation, Visualization, Writing – review & editing**. L.N.:** Formal analysis, Writing – review & editing**. C.C.I.:** Conceptualization, Funding acquisition, Project administration, Supervision, Writing – review & editing**. R.J.P.H.:** Conceptualization, Funding acquisition, Investigation, Methodology, Project administration, Resources, Supervision, Writing – review & editing.

## Acknowledgments

This work was funded by a Royal Society University Research Fellowship to R.J.P.H. (URF\R1\231537) and a Biotechnology and Biological Sciences Research Council grant to R.J.P.H. and C.C.I. (BB/V001388/1). K.D. was funded by R.J.P.H.’s URF and a BBSRC Fellowship (UKRI895). J.T. was funded by a BBSRC grant (BB/Y010892/1) and A.A. was funded by The Clarendon Fund in partnership with the Merton College Biology Scholarship.

