## Supplementary Materials for "DigiFish: A flexible open-source tool for animating realistic virtual fish and reconstructing visual fields from tracking data"

### DigiFish Supplementary Methods

To assess the similarity of DigiFish motion to that of a real fish, we compared the elementary motion detector (EMD) output of real zebrafish (*Danio rerio*) to that of digital twins constructed in DigiFish. EMDs, which are inspired by the motion-processing circuits of real animals, compare changes in luminance between video frames, producing estimates of the perceived magnitude and direction of motion in a scene (Santon et al., 2025).

We filmed two trials of real fish, each containing four adult zebrafish (two males and two females), in a circular arena (80 cm diameter) containing 1 cm of sand substrate, 6 cm of water above the sand, and five cones. The arena was illuminated from above with an LED light (NEEWER MS150B 130W Bi-Color LED Video Light setup) and a UV light with a high frequency ballast (18 Watt 24 Inch T8 UVA Tube Lamp).

We filmed the fish using four GoPro HERO13 cameras, each recording at 240 frames per second and 2.7K resolution. One camera was positioned directly above the centre of the arena, and three additional cameras were placed equidistantly around the arena's circumference, with views directed horizontally into the arena. These lateral cameras therefore captured views perpendicular to the overhead camera. Cameras were synchronised by flashing a light at the end of each trial.

We manually identified periods in which a single fish swam past a lateral camera in near isolation. The corresponding overhead footage was loaded into the AI-assisted tracking software SLEAP AI (version 1.4.1a2). Four anatomical landmarks were labelled on each fish: Head (tip of the snout), COM (centre of mass), Caudal (anterior caudal peduncle, at the posterior end of the dorsal fin), and Tail (distal end of the caudal peduncle). These points were manually annotated for every consecutive frame during which the target fish passed the lateral camera. The resulting tracks were exported from SLEAP AI as x-y coordinates and imported into DigiFish.

In DigiFish, we loaded a model zebrafish and dictated its movement using the tracking annotations from the real fish (with smoothing values set at 0.06). We then aligned the virtual lateral camera to match the position and orientation of the corresponding real lateral camera and adjusted lighting levels to approximate real-world conditions. Synthetic lateral frames were exported at the same resolution and frame rate as the real footage.

We processed the corresponding lateral frames from each dataset with an EMD model implemented in ImageJ (version 1.54g) using the following parameters: 3 cycles per degree (cpd) spatial frequency, 90° image angular width, and Gaussian sigma = 1.2. The resulting EMD output provides per-frame estimates of motion in the up, down, left, and right directions.
